# The roles of TGFβ and serotonin signaling in regulating proliferation of oocyte precursors and germline aging

**DOI:** 10.1101/2024.05.08.593208

**Authors:** Erin Z. Aprison, Svetlana Dzitoyeva, Ilya Ruvinsky

## Abstract

The decline of oocyte quality in aging but otherwise relatively healthy individuals compels a search for underlying mechanisms. Building upon a finding that exposure to male pheromone ascr#10 improves oocyte quality in *C. elegans*, we uncovered a regulatory cascade that promotes proliferation of oocyte precursors in adults and regulates oocyte quality. We found that the male pheromone promotes proliferation of oocyte precursors by upregulating LAG-2, a ligand of the Notch-like pathway in the germline stem cell niche. LAG-2 is upregulated by a TGFβ-like ligand DAF-7 revealing similarity of regulatory mechanisms that promote germline proliferation in adults and larvae. A serotonin circuit that also regulates food search and consumption upregulates DAF-7 specifically in adults. The serotonin/DAF-7 signaling promotes germline expansion to compensate for oocyte expenditure which is increased by the male pheromone. Finally, we show that the earliest events in reproductive aging may be due to declining expression of LAG-2 and DAF-7. Our findings highlight neuronal signals that promote germline proliferation in response to the environment and argue that deteriorating oocyte quality may be due to reduced neuronal expression of key germline regulators.

## Introduction

Loss of reproductive function is among the early detrimental effects of aging. For example, whereas life expectancy for women in the United States is ∼79 years (Xu et al., 2022), few women older than 45 have oocytes that could support embryonic development (Centers for Disease Control and Prevention, 2018). Similarly, *C. elegans* hermaphrodites become unable to produce fertilizable oocytes around mid-lifespan (Hughes et al., 2007). Reproductive aging in *C. elegans* hermaphrodites manifests as morphologically defective oocytes (Garigan et al., 2002), increased rates of aneuploidy (Luo et al., 2010), and depleted stores of mitotic germline precursors (Kocsisova et al., 2019; Qin and Hubbard, 2015), similar to observations in other species (Ishibashi et al., 2020; Miller et al., 2014; Nagaoka et al., 2012; Zhao et al., 2008). Discovery of mechanisms that maintain germline quality may lead to treatments that ameliorate the effects of reproductive aging (Achache et al., 2021; Templeman et al., 2018).

We previously reported (Aprison and Ruvinsky, 2016) that exposure of adult hermaphrodites to the major male pheromone ascr#10 (Izrayelit et al., 2012) slowed down the loss of germline precursors due to increased proliferation (Aprison et al., 2022a). Exposure during larval stages did not alter germline expansion (Aprison et al., 2022a). Hermaphrodites irreversibly switch to oocyte production late in larval development (Kimble and Crittenden, 2007); therefore, in adults, germline precursors are oocyte precursors. A consequence of greater supply of oocyte precursors in adults is increased physiological cell death in the germline, a process known to be required for oocyte quality maintenance (Andux and Ellis, 2008). Indeed, exposure to ascr#10 improves oocyte morphology and reduces aneuploidy and embryonic lethality of progeny (Aprison et al., 2022a). We wish to understand the mechanisms by which this male pheromone exerts salubrious effects on oocyte health. Here, we focused on neuronal regulation of germline events.

Development of the hermaphrodite germline in *C. elegans* has been extensively studied (Hubbard and Schedl, 2019). Germline proliferation in larvae is promoted by the Notch-like signaling pathway (Figure 1A) that principally consists of the ligand LAG-2 (Henderson et al., 1994) in the single-cell germline stem cell niche, the Distal Tip Cell (DTC), and the GLP-1 receptor in the germline (Crittenden et al., 1994). Downstream of GLP-1 signaling, an elaborate genetic network (Hubbard and Schedl, 2019) determines the balance between stem cell fate (i.e., proliferation) and meiotic entry (i.e., differentiation). We started our inquiry by testing the role of GLP-1 signaling in the effects of ascr#10 and continued by investigating the upstream neuronal signals that modulate this signaling in the adult hermaphrodite germline.

**Figure 1.**
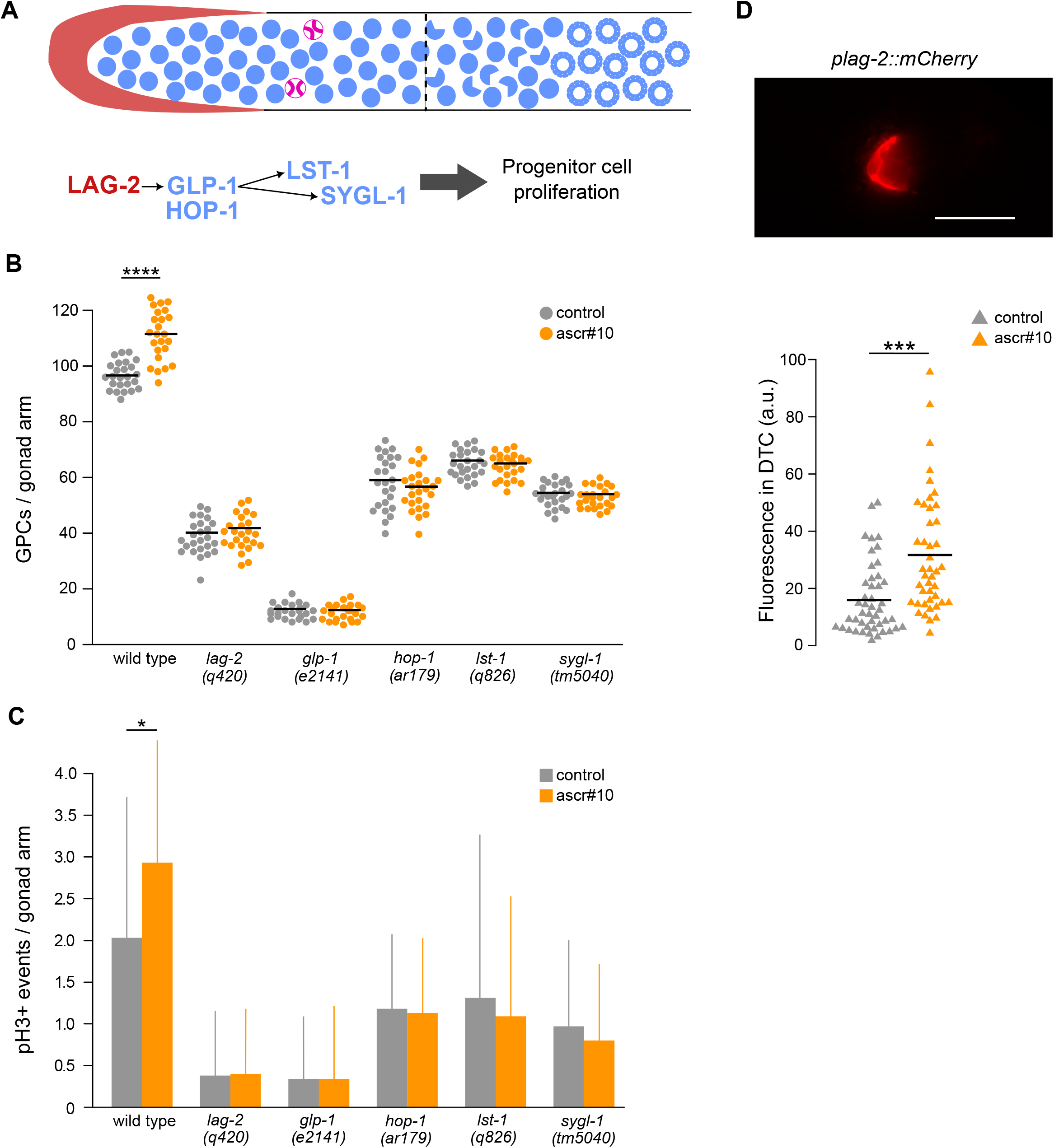
ascr#10 promotes germline proliferation in hermaphrodites via LAG-2/GLP-1 signaling. **(A)** Schematic depiction of the distal germline. The distal tip cell (DTC) is shown in red, germline cells are blue, and pH3+ cells are magenta. The dashed line denotes the boundary of the transition zone. A simplified LAG-2/GLP-1 signaling pathway is shown below. **(B)** GPC counts with or without exposure to ascr#10. Wild type, *hop-1, lst-1*, and *sygl-1* were scored on Day 2 of adulthood. *lag-2* and *glp-1* strains were shifted to non-permissive temperature as young adults and counted at the time equivalent to Day 2 of adulthood. Each dot represents a single gonad arm. Black bars are means. **(C)** Counts of pH3+ cells with or without exposure to ascr#10. Worms were treated and scored as in panel B. Error bars are standard deviation. **(D)** A representative image of DTC expression of *plag-2::mCherry*; quantification shown below. Each triangle is the measurement of one DTC; black bars denote means. Scale bar = 20μm. * p<0.05, *** p<0.001, **** p<0.0001. Additional data in Figure S1. See Table S1 for sample sizes and statistical analyses.

## Results & Discussion

### ascr#10 promotes germline proliferation in hermaphrodites via LAG-2/GLP-1 signaling

We tested loss-of-function mutations in genes encoding the ligand (*lag-2*) and the receptor (*glp-1*) of the Notch-like signaling in the germline. We also tested two direct downstream targets of GLP-1 signaling, SYGL-1 and LST-1 (Chen et al., 2020; Kershner et al., 2014; Shin et al., 2017), and because ascr#10 effects on germline proliferation are restricted to adults (Aprison et al., 2022a; Aprison and Ruvinsky, 2019b), the adult-specific presenilin HOP-1 (Agarwal et al., 2018). Mutations in all five of these genes prevented the ascr#10-dependent increase of the number of germline precursor cells (GPCs) (Figure 1B) likely due to the failure to increase germline proliferation (Figure 1C).

Because ascr#10 promotes germline proliferation (Aprison et al., 2022a) and because the GLP-1 pathway is required for the ascr#10-dependent increase in proliferation, we tested whether the pheromone upregulates expression of the ligand LAG-2. We found that expression of both a *lag-2* promoter-driven transgene (Pekar et al., 2017) (Figure 1D) and an endogenously tagged LAG-2::mNeonGreen protein (Gordon et al., 2019) (Figure S1) was significantly higher in the presence of ascr#10. We concluded that ascr#10 promotes germline proliferation in hermaphrodites by increasing GLP-1 signaling due to transcriptional upregulation of the LAG-2 ligand.

### Serotonin signaling increases proliferation in adult hermaphrodite germlines

Next, we sought to identify upstream regulators of LAG-2. In a series of prior studies, we documented the role of signaling via a specific serotonergic circuit in increasing germline proliferation in response to ascr#10. Exposure to ascr#10 increases serotonin signaling from HSN and NSM neurons (Aprison and Ruvinsky, 2019a), although only in egg-laying adults (Aprison and Ruvinsky, 2019b), to the MOD-1 receptor in a small subset of neurons (Aprison and Ruvinsky, 2019a; Aprison and Ruvinsky, 2022b).

Most relevant to the present study, we showed that serotonin signaling is required for ascr#10 to improve oocyte quality (Aprison et al., 2022b) and that stimulation of serotonin signaling with low doses of serotonin reuptake inhibitors increases germline proliferation and improves oocyte quality (Aprison et al., 2023). We therefore tested whether the same pharmacological stimulation caused transcriptional upregulation of LAG-2. It did (Figure 2A).

**Figure 2.**
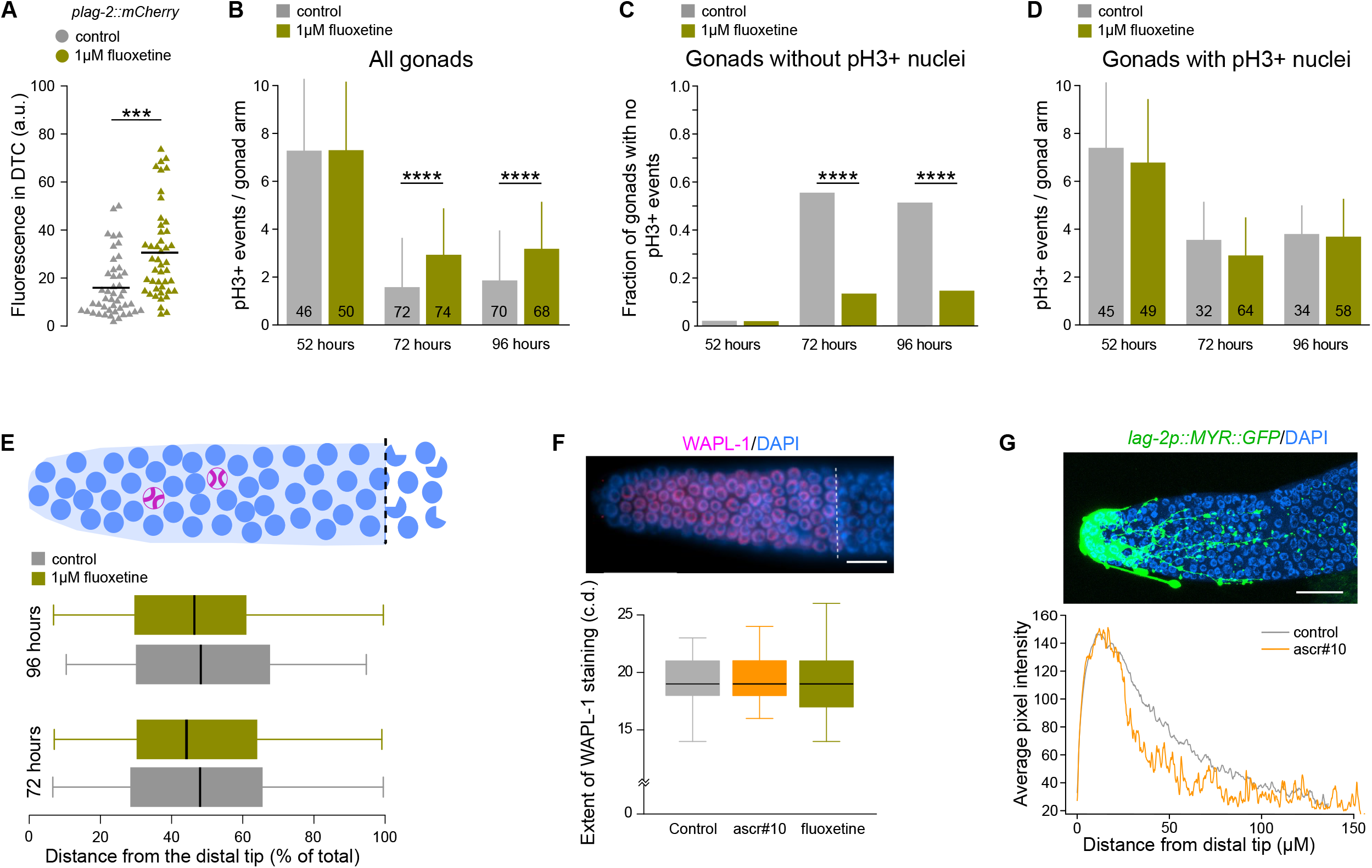
Serotonin signaling increases germline proliferation in adult hermaphrodites. **(A)** The effect of fluoxetine on expression of *plag-2::mCherry* in hermaphrodite DTC. Each triangle represents one DTC; black bars denote means. **(B)** pH3+ events in hermaphrodites treated with fluoxetine and control measured at 52 hours (young adults prior to the onset of egg laying), 72 hours (Day 2 adults), and 96 hours (Day 3 adults). Numbers in bars show the numbers of gonads examined. **(C)** Fraction of the total gonad arms counted that had no pH3+ events. **(D)** Numbers of pH3+ events in the gonads of hermaphrodites with at least one pH3+ event. Numbers in bars show the number of gonads counted. In **B** and **D**, error bars are standard deviations. **(E)** Positions of pH3+ events as measured as a percentage of the distance between the DTC (left) and the transition zone (right). Schematic above as in Figure 1A. Black lines are medians, boxes represent the two middle quartiles, whiskers 95%. **(F)** A representative image and quantification of the extent (in cell diameters, c.d.) of anti-WAPL-1 antibody staining (magenta) against the background of DAPI-stained cells (blue). The dashed line denotes the boundary of the transition zone. The meaning of elements of the box plots as in **E. (G)** A representative image of the DTC (green) in a *lag-2p::MYR::GFP* hermaphrodite; DAPI-stained germline cells are blue. Quantification of plot profiles of GFP fluorescence taken from the average of maximum projection confocal images. *** p<0.001, **** p<0.0001. Error bars are standard deviation. Scale bars = 10μm. Additional data in Figure S2. See Table S1 for sample sizes and statistical analyses.

Consistent with our previous findings (Aprison et al., 2022a; Aprison et al., 2023; Aprison and Ruvinsky, 2019b), prior to the onset of egg laying, fluoxetine had no discernable effect on germline proliferation (Figure 2B). Adults have reduced germline proliferation compared to late larvae (Roy et al., 2016). We found that even though there were fewer proliferative events in the gonads of Day 2 and 3 adults than in those immediately after the L4-to-adult molt, fluoxetine significantly boosted proliferation (Figure 2B). It was noted previously that many gonad arms in aging adult hermaphrodites have zero M-phase nuclei (Cinquin et al., 2016; Maciejowski et al., 2006). Fluoxetine reduced the fraction of such gonad arms by nearly five-fold (Figure 2C), while the average number of divisions in the germlines with at least one division was essentially the same between control and treatment populations (Figure 2D).

We found no discernable differences in spatial distributions of division events throughout the distal germline (Figure 2E) and the location of the transition zone (Figure 2F, Figure S2A) between control and fluoxetine-treated animals. Likewise, we did not observe reproducible differences in the morphology or the extent of elaboration of the DTC using transgenic markers (Byrd et al., 2014; Gordon et al., 2020) (Figure 2G, Figure S2B). We concluded that increased serotonin signaling upregulates LAG-2 expression in the DTC of adult hermaphrodites consequently reducing the number of germline arms without division events.

### Serotonin acts upstream of DAF-7 to regulate germline proliferation in adult hermaphrodites

While we demonstrated that serotonin regulates LAG-2 expression in adults, prior work showed that in larvae this ligand of the Notch pathway is non-cell-autonomously regulated by DAF-7, a member of the TGFβ family (Dalfo et al., 2012; Pekar et al., 2017). DAF-7 is required for the increase in the number of GPCs upon exposure to ascr#10 (Aprison and Ruvinsky, 2017). To clarify the relationships between the serotonin and DAF-7 signaling pathways with germline proliferation in adults we carried out additional experiments.

We found that transcriptional upregulation of LAG-2 in the DTC by ascr#10 was strictly dependent on DAF-7 (Figure 3A), as was the increase in the number of GPCs (Figure 3B). These results implied that LAG-2 upregulation mediated by DAF-7 is the cause of increased germline proliferation and greater number of GPCs in the presence of ascr#10. Expression of DAF-7 in ASI neurons (we did not observe consistent expression of the transcriptional reporter elsewhere) was likewise increased by ascr#10 (Figure 3C). Because in all our experiments fluoxetine (i.e., increased serotonin signaling) phenocopied the effects of ascr#10 (Figure 3A-C), we hypothesized that serotonin acts upstream of DAF-7. Confirming this inference, loss of TPH-1 (an enzyme essential for serotonin production (Sze et al., 2000)) prevented transcriptional upregulation of DAF-7 expression in ASI neurons by ascr#10 (Figure 3D).

**Figure 3.**
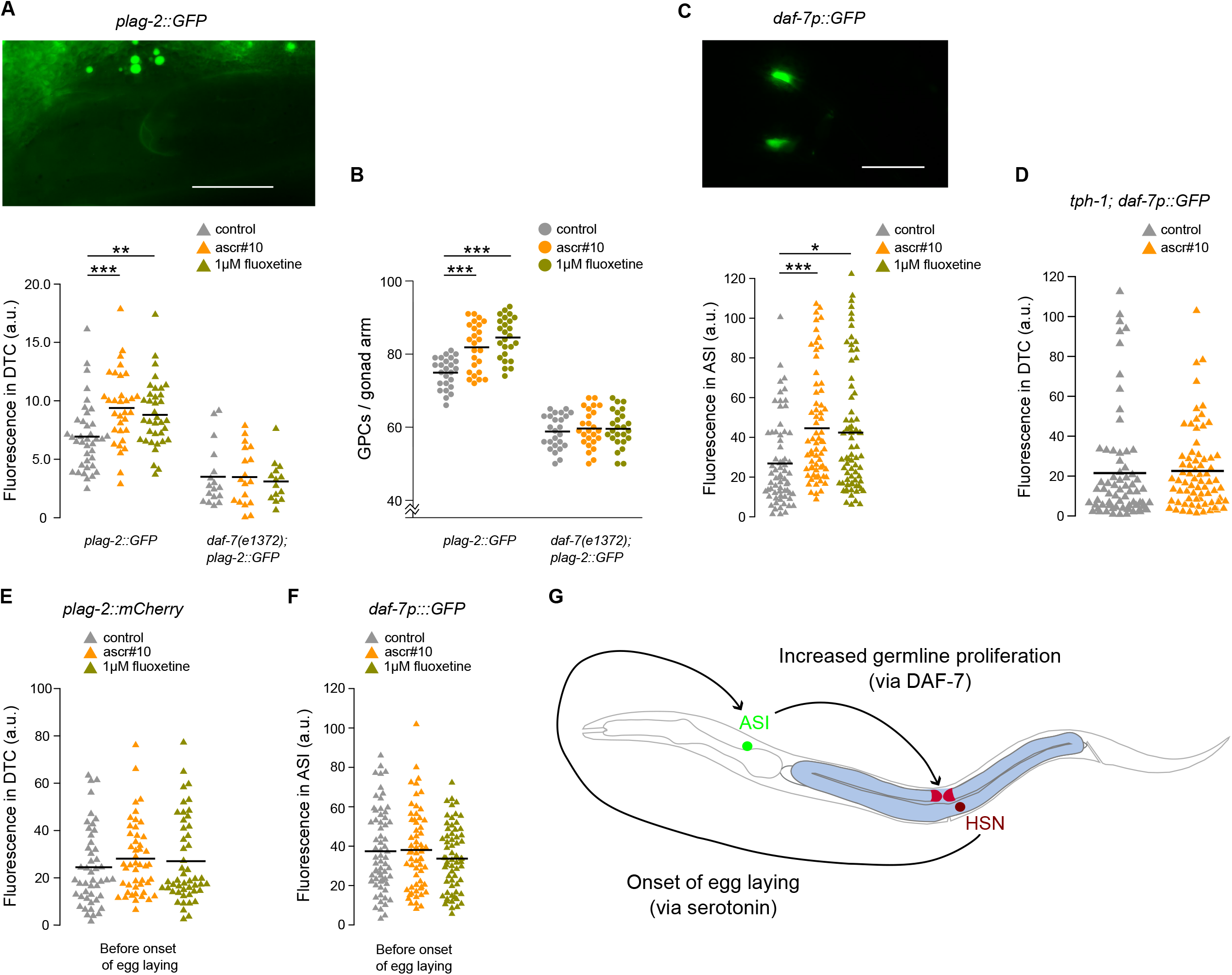
Serotonin acts upstream of DAF-7 to regulate germline proliferation in adult hermaphrodites. **(A)** A representative image of *plag-2::GFP* expression in the DTC. Below is the quantification of fluorescence in the DTC. Worms were raised at 15°C, transferred to 20°C at 96 hours, and imaged at 120 hours (the equivalent of young adults after the onset of egg laying). **(B)** GPC counts. **(C)** Representative image of *daf-7p::GFP* fluorescence in ASI neurons. Quantification of fluorescence in the ASI neurons in hermaphrodites after the onset of egg laying. **(D)** *daf-7p::GFP* fluorescence in the ASI neurons of *tph-1* loss-of-function mutants after the onset of egg laying. **(E)** *plag-2::mCherry* fluorescence in the DTC of young adults before the onset egg laying. **(F)** *daf-7p::GFP* fluorescence in the ASI neurons of young adults before the onset of egg laying. In **A – F**, black bars are means; triangles represent one DTC (**A, D, E**) or one ASI neuron (**C, F**); circles one gonad arm (**B**). **(G)** A model of neuronal signaling that promotes germline proliferation in response to increased egg laying. *p<0.05, ** p<0.01, *** p<0.001. Scale bars = 20μm. Additional data in Figure S3. See Table S1 for sample sizes and statistical analyses.

We concluded that in adult hermaphrodites upregulated DAF-7 expression in ASI neurons increases LAG-2 expression in the DTC and thus stronger activation of Notch/GLP-1 signaling and proliferation of germline precursors. The same regulatory relationship appears to operate during larval expansion of the germline – mutations that reduce activity of the TGFβ/DAF-7 pathway have fewer germline precursors (Dalfo et al., 2012; Pekar et al., 2017).

The one nuance that differentiates our study from those of Dalfo et al. and Pekar et al. is that whereas we saw increased germline proliferation on “pheromone”, they observed decreased proliferation. We believe this difference is explained by the fact that whereas we studied the male pheromone ascr#10, Dalfo et al. and Pekar et al. tested overall dauer pheromone, a blend of several molecules that contains ascr#3 (Srinivasan et al., 2008). ascr#10 and ascr#3 generally have opposing effects on hermaphrodites (Aprison and Ruvinsky, 2015). Specifically, while ascr#10 promotes germline proliferation, ascr#3 counters this effect (Aprison and Ruvinsky, 2017; Aprison and Ruvinsky, 2022a).

### Egg laying licenses DAF-7 and LAG-2 responses to ascr#10

The *C. elegans* egg-laying circuit has feedforward and feedback features. The command motoneuron, HSN (Schafer, 2006), initiates episodes of egg laying by releasing serotonin and neuropeptides (Brewer et al., 2019). Mechanical deformations caused by the passage of the egg through the vulva initiates feedback from muscle and neuroendocrine cells onto the HSN (Collins et al., 2016; Medrano and Collins, 2023; Ravi et al., 2018; Yan et al., 2024). In response to feedback from the vulva, HSN signals to modify behavioral programs related to search and consumption of food (Huang et al., 2023; Ravi et al., 2019).

We have previously identified elements of the circuit that limits to egg-laying adults the increase of germline proliferation in response to ascr#10 (Aprison et al., 2022a; Aprison and Ruvinsky, 2019b) and fluoxetine (Aprison et al., 2023). Upon exposure to the pheromone, increased signaling from serotonin-producing neurons HSN and pharyngeal NSM inhibits a small subset of target neurons via the MOD-1 receptor (Aprison and Ruvinsky, 2019a; Aprison and Ruvinsky, 2022b). The signal from NSM/HSN is counteracted by serotonin transporter MOD-5 that acts in serotonin-absorbing neurons AIM and RIH (Aprison et al., 2022b). However, in the absence of feedback signaling from the vulva that communicates mechanical distortion of passing of embryos, ascr#10 does not increase serotonin signaling (Aprison and Ruvinsky, 2019b). This organization of the circuit limits ascr#10 responses – germline proliferation and behavioral – to actively reproducing animals.

ascr#10 promotes egg laying and mating receptivity (Aprison et al., 2022b; Aprison and Ruvinsky, 2019a) as well as reduces exploration (Aprison and Ruvinsky, 2019a) and increases pharyngeal pumping (Angeles-Albores et al., 2023). The coupling between reproductive and food seeking behaviors likely assures that sufficient resources are consumed to match the increased reproductive output. Similarly, the regulation of germline proliferation by the same serotonin circuit increases production of germline precursors in the presence of a pheromone that increases the rate of egg laying.

We further investigated how this serotonin-mediated feedback from egg laying regulates germline proliferation. First, we established that adult hermaphrodites that have not yet started laying eggs did not increase expression of LAG-2 in response to either ascr#10 or fluoxetine (Figure 3E, Figure S3), unlike their slightly older (∼4-6 hours) counterparts that have started laying eggs (Figure 3A). Second, prior to onset of egg laying neither ascr#10 nor increased serotonin signaling increased DAF-7 expression in ASI neurons (Figure 3F). Combined with our previous demonstration that ascr#10 does not increase serotonin signaling prior to onset of egg laying (Aprison and Ruvinsky, 2019b), these results suggest a simple model. After the onset of egg laying that licenses activity of the circuit, ascr#10 increases serotonin signaling via the HSN-containing circuit to upregulate DAF-7 in ASI neurons, which non-cell-autonomously causes greater production of LAG-2 in the DTC, thereby increasing GLP-1/Notch signaling and germline proliferation (Figure 3G). In so doing, the serotonin/DAF-7/LAG-2 signaling increases production of oocyte precursors to replenish mature oocytes that are expended at a greater rate in the presence of ascr#10. The totality of behavioral and developmental changes caused by ascr#10 improves reproductive success on this pheromone (Aprison and Ruvinsky, 2019a).

### Reduced TGFβ signaling as a driver of germline aging

As reported previously (Roy et al., 2016), adult hermaphrodites have fewer proliferating germline precursors than late larvae (Figure 2B). Comparing young adults just prior to egg laying, slightly older adults that started laying eggs, and Day 2 adults suggested that progressive reduction of expression levels of LAG-2 in the DTC was the likely proximal cause for less proliferation in adults (Figure 4A). Exposure to ascr#10 or fluoxetine increased LAG-2 expression in the egg-laying animals to the levels comparable to those in late L4/early pre-reproductive adults (Figure 4B). Similarly, expression of DAF-7 was significantly lower in egg-laying adults than in animals that are just a few hours younger (Figure 4C), while ascr#10 and fluoxetine boosted it to the late larval level (Figure 4D).

**Figure 4.**
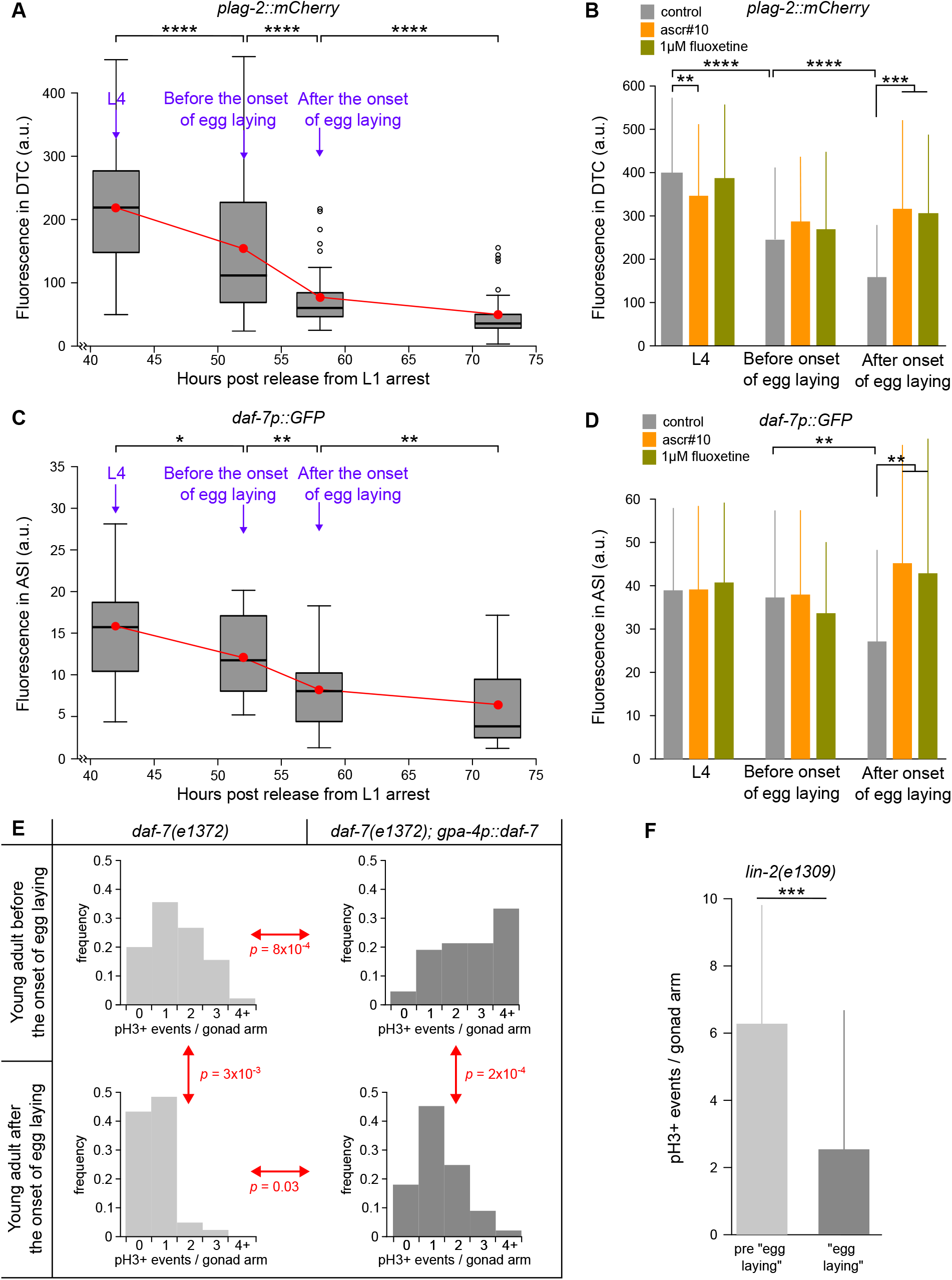
Reduced neuronal TGFβ signaling as a driver of germline aging. **(A)** Quantification of *plag-2::mCherry* fluorescence in the DTC. **(B)** Quantification of *plag-2::mCherry* fluorescence in the DTC in hermaphrodites treated with ascr#10 or fluoxetine. **(C)** Quantification of *daf-7p::GFP* fluorescence in ASI neurons. **(D)** Quantification of *daf-7p::GFP* fluorescence in ASI neurons in hermaphrodites treated with ascr#10 or fluoxetine. In **A** and **C**, black bars represent medians, red dots represent means. In **B** and **D**, error bars are standard deviation. **(E)** Distributions of numbers of pH3+ events per gonad arm in *daf-7(e1372)* and *daf-7(e1372); gpa-4p::daf-7* hermaphrodites before or after onset of egg laying. **(F)** pH3+ events in the vulvaless *lin-2(e1309)* hermaphrodites. Worms were judged to be “pre egg laying” or “egg laying” based on whether wild type worms would have initiated egg laying having as many embryos in the uterus (<4 and >6 embryos, respectively). Error bars are standard deviation. * p<0.05, ** p<0.01, *** p<0.001, **** p<0.0001. See Table S1 for sample sizes and statistical analyses.

We sought further support for the idea that age-dependent decline in DAF-7 signaling causes progressively reduced germline proliferation in older hermaphrodites. We found that constitutive expression (You et al., 2008) of DAF-7 in ASI (and AWA (Kim et al., 2005)) neurons significantly increased germline proliferation while nearly eliminating gonadal arms without division events (Figure 4E). While the onset of egg laying is a watershed moment in regulation of germline proliferation (Figure 2B, 2C; Figure 3A vs. Figure 3E; Figure 3C vs. Figure 3F), it is not solely responsible for reduced cell division in adults – the drop is observed even in vulvaless mutants (Figure 4F). Moreover, the decline of DAF-7 and LAG-2 expression is evident both before and after the onset of egg laying (Figure 4A, Figure 4C).

These results have implications for understanding germline aging. An early sign of germline aging in *C. elegans* hermaphrodites is the reduced number of germline stem cells (Luo et al., 2010; Qin and Hubbard, 2015) due to lower proliferation (Achache et al., 2021) likely caused by reduced expression of components of the Notch/GLP-1 signaling (Kocsisova et al., 2019). Our findings are consistent with this view (Figure 4A). Some aspects of germline aging are delayed by overexpressing SYGL-1, a target of GLP-1 signaling, in the distal gonad (Kocsisova et al., 2022). The most plausible interpretation of improved oocyte quality in hermaphrodites in response to ascr#10 (Aprison et al., 2022a) or serotonin reuptake inhibitors (Aprison et al., 2023) is that both of these treatments restore youthful levels of neuronal DAF-7 signaling, which causes increased LAG-2/GLP-1 signaling and greater germline proliferation. In this paradigm, progressively declining DAF-7 signaling non-cell-autonomously drives germline aging. Factors causing the decline of DAF-7 are not currently known, but are unlikely to be the onset of egg laying (Figure 4C) or serotonin decline (expression of TPH-1 is relatively steady during early adulthood (Aprison and Ruvinsky, 2019b)). We propose that decreasing signals from other types of neurons contribute to the onset and progression of germline aging.

## Acknowledgements

We are grateful to Rick Morimoto for generous hospitality. We thank Frank Schroeder for the gift of ascr#10, Kacy Gordon, Jane Hubbard, and Tim Schedl for sharing strains, Sarah Crittenden for advice, and Kacy Gordon for thoughtful comments on this project and the manuscript. This work was funded in part by NIH (R01GM126125) grant to IR. We thank WormBase and the Caenorhabditis Genetics Center (CGC). WormBase is supported by grant U41 HG002223 from the National Human Genome Research Institute at the NIH, the UK Medical Research Council, and the UK Biotechnology and Biological Sciences Research Council. The CGC is funded by the NIH Office of Research Infrastructure Programs (P40 OD010440).

## Materials and Methods

### *C. elegans* handling and strains

*C. elegans* nematodes were maintained under standard conditions at 20°C as described previously (Aprison et al., 2022a; Aprison et al., 2023; Aprison and Ruvinsky, 2017; Aprison and Ruvinsky, 2019a; Aprison and Ruvinsky, 2019b). A complete strain list is included in Table S1. Synchronization was done by hypochlorite treatment and age was measured as time from release from the L1 arrest as outlined earlier (Aprison et al., 2022a; Aprison et al., 2023; Aprison and Ruvinsky, 2017; Aprison and Ruvinsky, 2019a; Aprison and Ruvinsky, 2019b). Strains that contained temperature sensitive alleles were maintained at 15°C and shifted to non-permissive temperature just after the L4/young adult molt except for *lin-2(e1309)* which was shifted to the non-permissive temperature after hatching. Plates were conditioned with ascr#10 (Aprison et al., 2022a; Aprison and Ruvinsky, 2017; Aprison and Ruvinsky, 2019a; Aprison and Ruvinsky, 2019b) and fluoxetine (Aprison et al., 2023) as described previously.

### Imaging

For counting GPCs, worms were DAPI-stained, mounted on agarose pads, and imaged precisely as described previously (Aprison et al., 2022a; Aprison and Ruvinsky, 2016; Aprison and Ruvinsky, 2017; Aprison and Ruvinsky, 2019a; Aprison and Ruvinsky, 2019b). For counting proliferative events, worms were dissected and stained with pH3 antibodies and imaged as described (Aprison et al., 2022a; Aprison et al., 2022b). Worms stained with anti-WAPL-1 were dissected and stained in the same manner. Strains JK4475 and NK2571 were imaged with a Zeiss LSM 800 confocal microscope. Fluorescence quantification and plot profile analysis were done using ImageJ.

### Statistical analyses

Tests of statistical significance were carried out in R and Excel. Sample sizes and p-values are shown in Table S1.

