## Supplemental Figures w legends for "The roles of TGFβ and serotonin signaling in regulating proliferation of oocyte precursors and germline aging"

*lag-2(cp193[lag-2::mNeonGreen^3xFlag])*

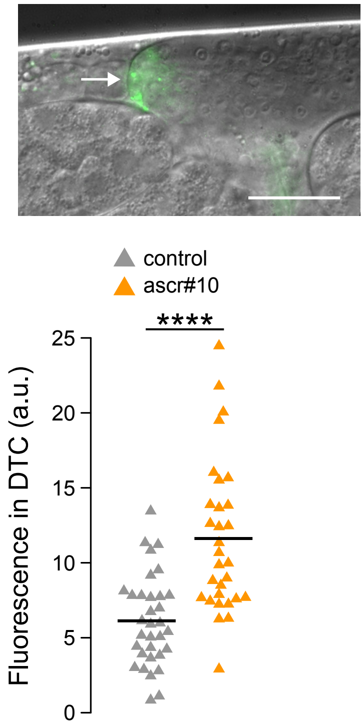

**Figure S1. ascr#10 increases the amount of LAG-2 protein in the DTC.**

A Nomarski image of the distal tip of one gonad arm overlaid with a false-colored image of endogenously tagged LAG-2 protein fused to mNeonGreen (Gordon et al., 2019). The arrow points to the location of the DTC. Scale bar = 20μm. LAG-2 protein accumulates in a punctate pattern in the DTC. Quantification of the protein is shown below. Each triangle represents the measurement of fluorescence in a single DTC. Black bars are means. ****p<0.0001. See Table S1 for sample sizes and statistical analyses.

**Gordon, K. L., Payne, S. G., Linden-High, L. M., Pani, A. M., Goldstein, B., Hubbard, E. J. A. and Sherwood, D. R.** (2019). Ectopic Germ Cells Can Induce Niche-like Enwrapment by Neighboring Body Wall Muscle. *Curr Biol* **29**, 823-833 e825.

**A**

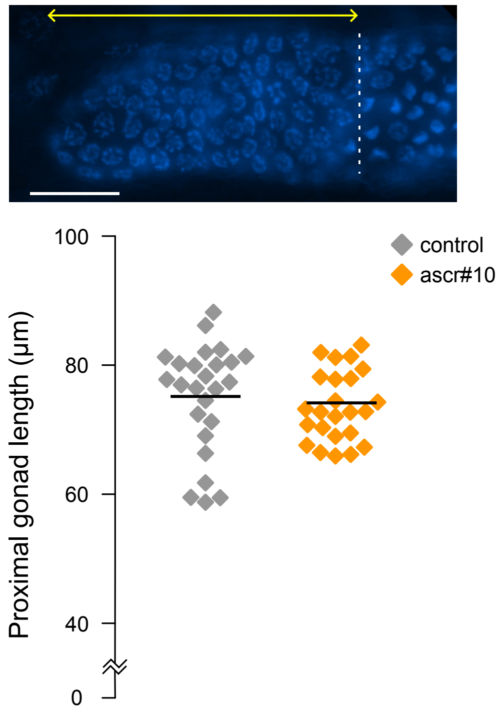


**B**

*cpIs122(lag-2p::mNeonGreen:: PLCδPH)*

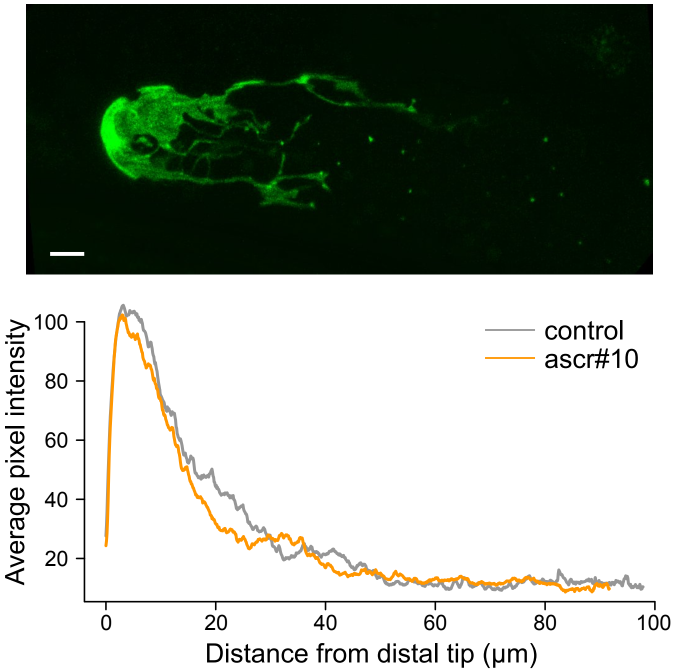

**Figure S2. ascr#10 does not change the length of the progenitor zone or the morphology of the DTC.**

**(A)** An image of a DAPI-stained gonad arm. Scale bar = 20μm. The dashed line denotes the boundary of the transition zone. The yellow double arrow spans the distance between the transition zone boundary and the distal tip of the gonad. These distances are not significantly different between ascr#10-treated and control hermaphrodites (each diamond = represents one gonad arm). **(B)** Maximum intensity projection of the gonad arm in a *cpIs122(lag-2p::mNeonGreen::PLCδPH)* worm (Gordon et al., 2020). Distal is to the left. Scale bar = 20μm. Below is the average plot profile of 10 control and 11 ascr#10 confocal experiments. See Table S1 for sample sizes and statistical analyses.

**Gordon, K. L., Zussman, J. W., Li, X., Miller, C. and Sherwood, D. R.** (2020). Stem cell niche exit in C. elegans via orientation and segregation of daughter cells by a cryptic cell outside the niche. *Elife* **9**.

*lag-2(cp193[lag-2::mNeonGreen^3xFlag])*

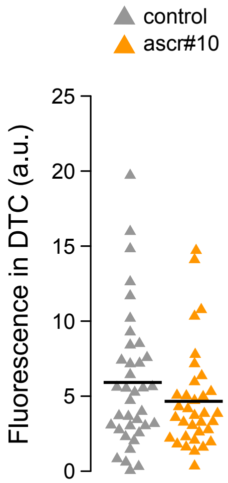

**Figure S3. ascr#10 does not increase the amount of LAG-2 protein in the DTC in worms before the onset of egg laying.**

Quantification of the accumulation of LAG-2 protein (fused to mNeonGreen (Gordon et al., 2019)) in young adult hermaphrodites before the onset of egg laying. Each triangle represents the measurement of fluorescence in a single DTC. Black bars are means. See Table S1 for sample sizes and statistical analyses.

**Gordon, K. L., Payne, S. G., Linden-High, L. M., Pani, A. M., Goldstein, B., Hubbard, E. J. A. and Sherwood, D. R.** (2019). Ectopic Germ Cells Can Induce Niche-like Enwrapment by Neighboring Body Wall Muscle. *Curr Biol* **29**, 823-833 e825.
